## Supplementary Material for "Modelling the synergistic effect of bacteriophage and antibiotics on bacteria: killers and drivers of resistance evolution"

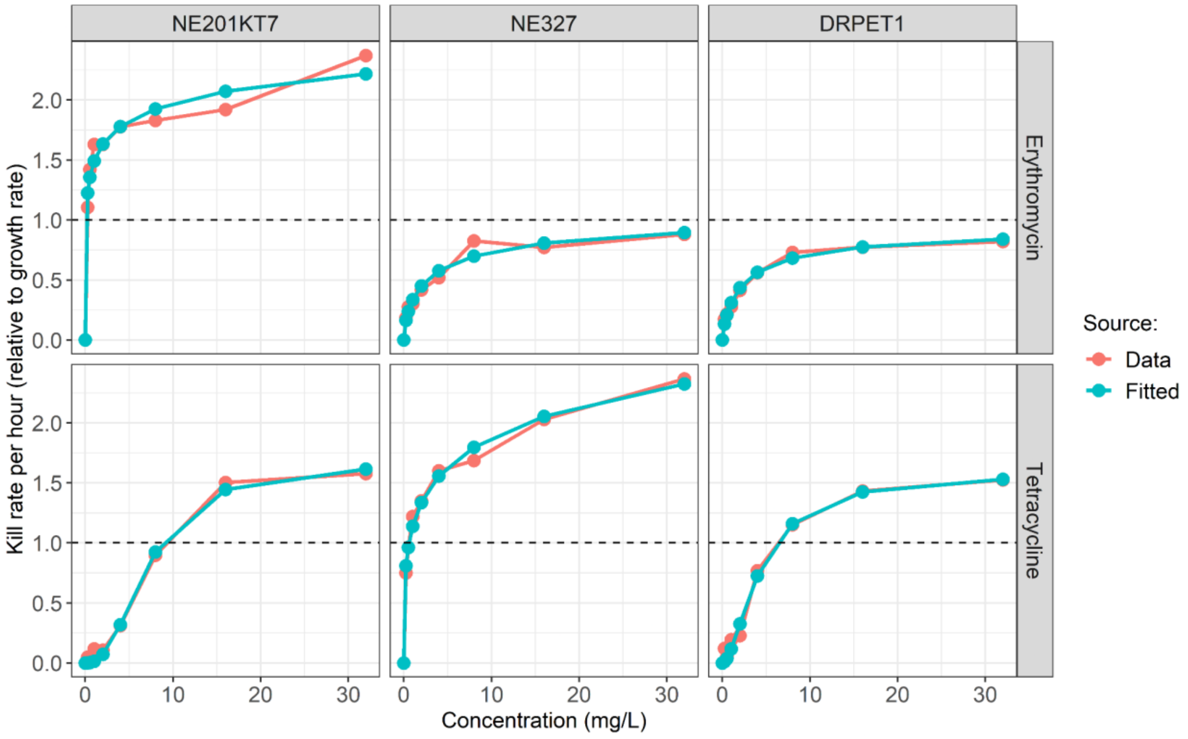


**Figure S1: Antibacterial effect of erythromycin and tetracycline measured *in vitro* (pink) and obtained after fitting Hill equations (blue).** Effect is relative to bacterial growth, such that a value greater than 1 indicates killing (net negative growth), while a value between 0 and 1 indicates only a decrease in growth rate. NE201KT7 contains a tetracycline-resistance gene (*tetK*), NE327 contains an erythromycin-resistance gene (*ermB*) and DRPET1 contains both resistance genes. The Hill equation is shown in Equation 3.

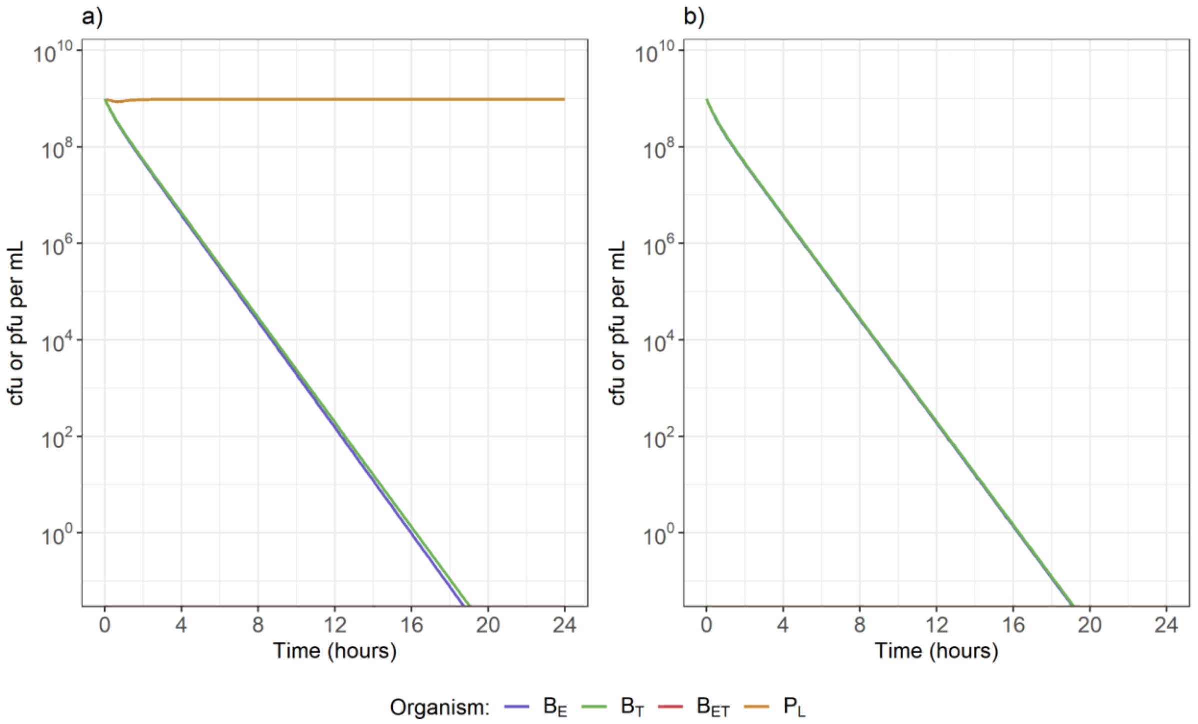


**Figure S2: a) The antibacterial effect of 1 mg/L of both erythromycin and tetracycline alongside $10^9$ pfu/mL of phage is equivalent to b) the effect of 4.58 mg/L of erythromycin and 1.14 mg/L of tetracycline in the absence of phage.** This was estimated by setting the concentration of phage to 0 in b) and fitting the concentrations of erythromycin and tetracycline to reproduce the decrease in bacteria numbers seen in a). cfu: colony-forming units; pfu: plaque-forming units.

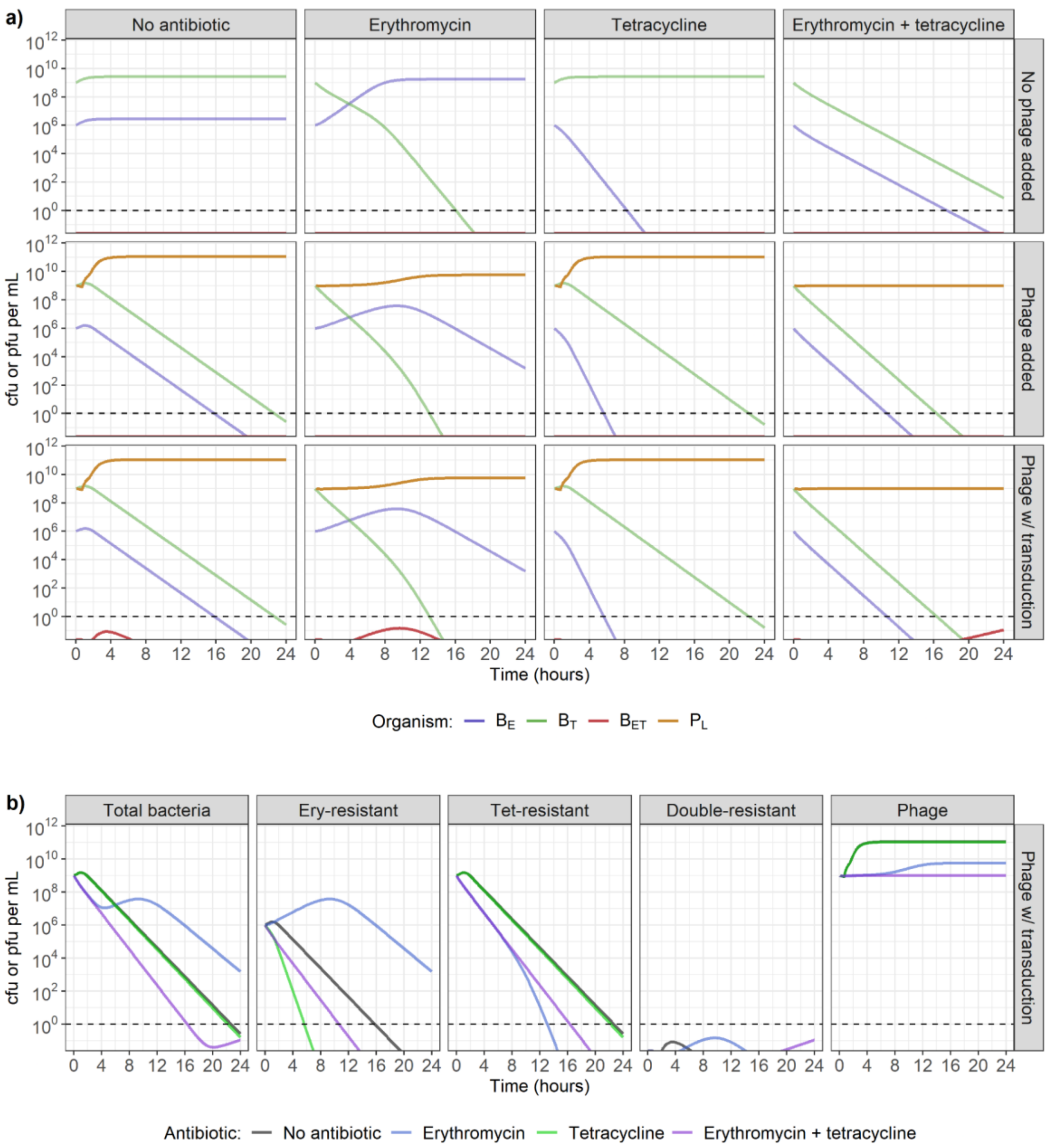


**Figure S3: a) Model-predicted bacterial dynamics in the presence of no antibiotics (1st column), erythromycin only (2nd column), tetracycline only (3rd column), or both erythromycin and tetracycline (4th column), combined with either no phage (top row), phage incapable of transduction (middle row), or phage capable of generalised transduction (bottom row).** Tetracycline-resistant bacteria ($B_T$) are initially present at a concentration of $10^9$ colony-forming units (cfu)/mL, and erythromycin-resistant bacteria ($B_E$) at $10^6$ cfu/mL. Antibiotics and/or phage ($P_L$) are present at the start of the simulation, at concentrations of 1 mg/L and $10^9$ plaque-forming units (pfu)/mL respectively. Double-resistant bacteria ($B_{ET}$) can be generated by generalised transduction only. Dashed line indicates the detection threshold of 1 cfu or pfu/mL. **b) Change in bacteria (single-resistant to erythromycin, single-resistant to tetracycline, or double-resistant) and phage numbers depending on the antibiotic exposure, in the presence of phage capable of generalised transduction.**

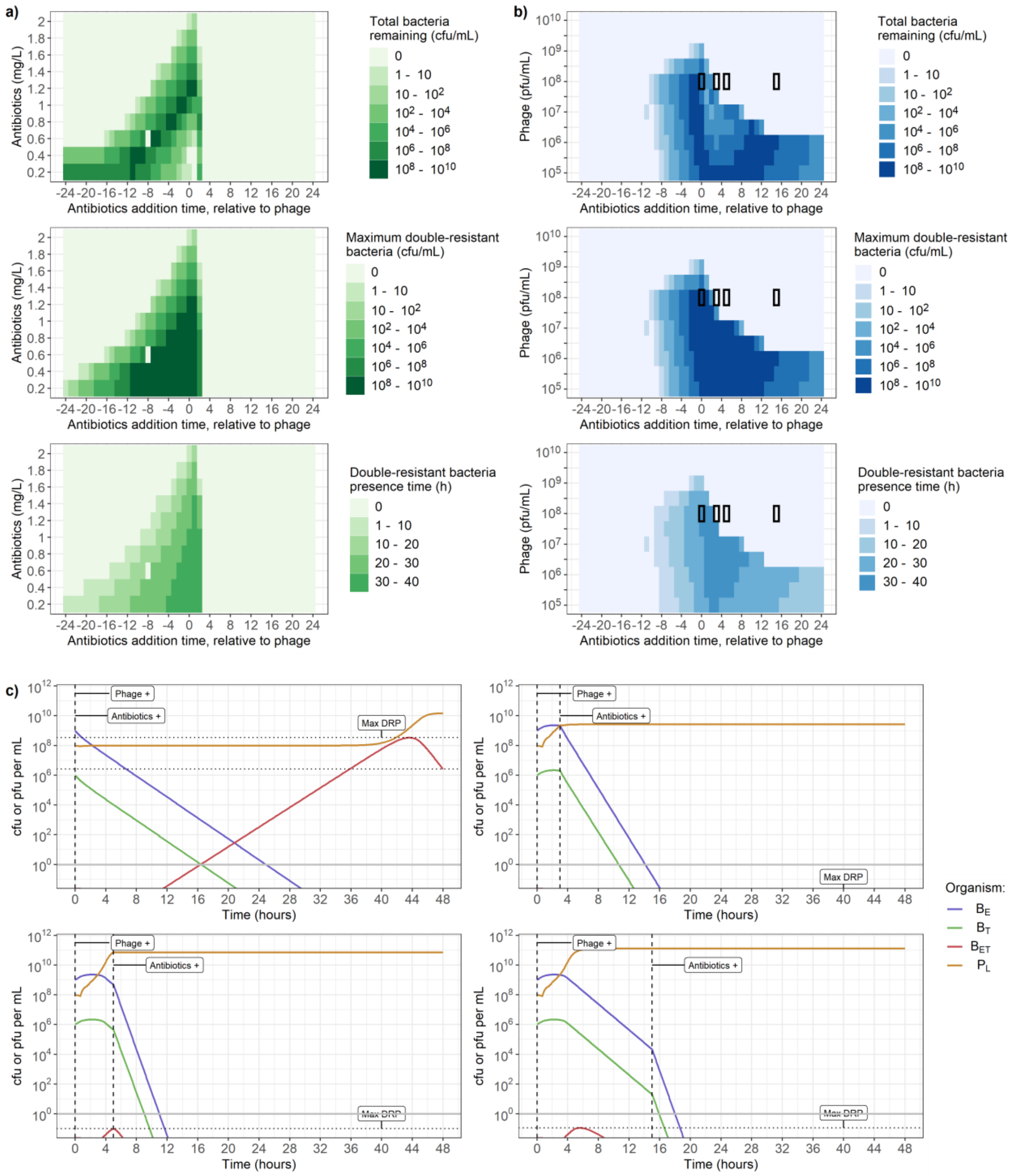


**Figure S4: Effect of varying antibiotic and phage timing and concentration when the tetracycline-resistant bacterial strain ($B_T$) is in minority ($10^6$ cfu/mL). a-b) Varying timing (x-axis) and dose of antibiotic and phage (y-axis) affects total bacterial count after 48h (top), maximum concentration of double-resistant bacteria ($B_{ET}$) (middle), and time when the concentration of $B_{ET}$ is greater than 1 colony-forming unit (cfu) per mL (bottom). a)** Adding $10^8$ plaque-forming units (pfu) per mL of phage, and between 0.2 and 2.2 mg/L of both erythromycin and tetracycline. **b)** Adding 1 mg/L of both erythromycin and tetracycline, and between $10^5$ and $10^{10}$ pfu/mL of phage. The x-axis indicates the time when antibiotics were added, relative to when phage were added. For example, the value "4"

indicates that phage were present at the start of the simulation, and antibiotics were introduced 4h later. The segments with black borders correspond to the dynamics shown in c). **c) Phage and bacteria dynamics over 48h for 4 conditions taken from panel b.** In all 4 conditions, indicated by the black rectangles, phage are initially present at a concentration of $10^8$ pfu/mL, while erythromycin and tetracycline are both introduced at concentrations of 1 mg/L after either 0h, 3h, 5h or 15h, stated on the plots, with the timing indicated by the vertical dashed lines. Horizontal dotted lines indicate bacteria remaining after 48h (corresponding to the top row of a-b) and maximum double-resistant bacteria ($B_{ET}$) concentration (middle row of a-b). Solid line indicates the detection threshold of 1 cfu or pfu/mL.

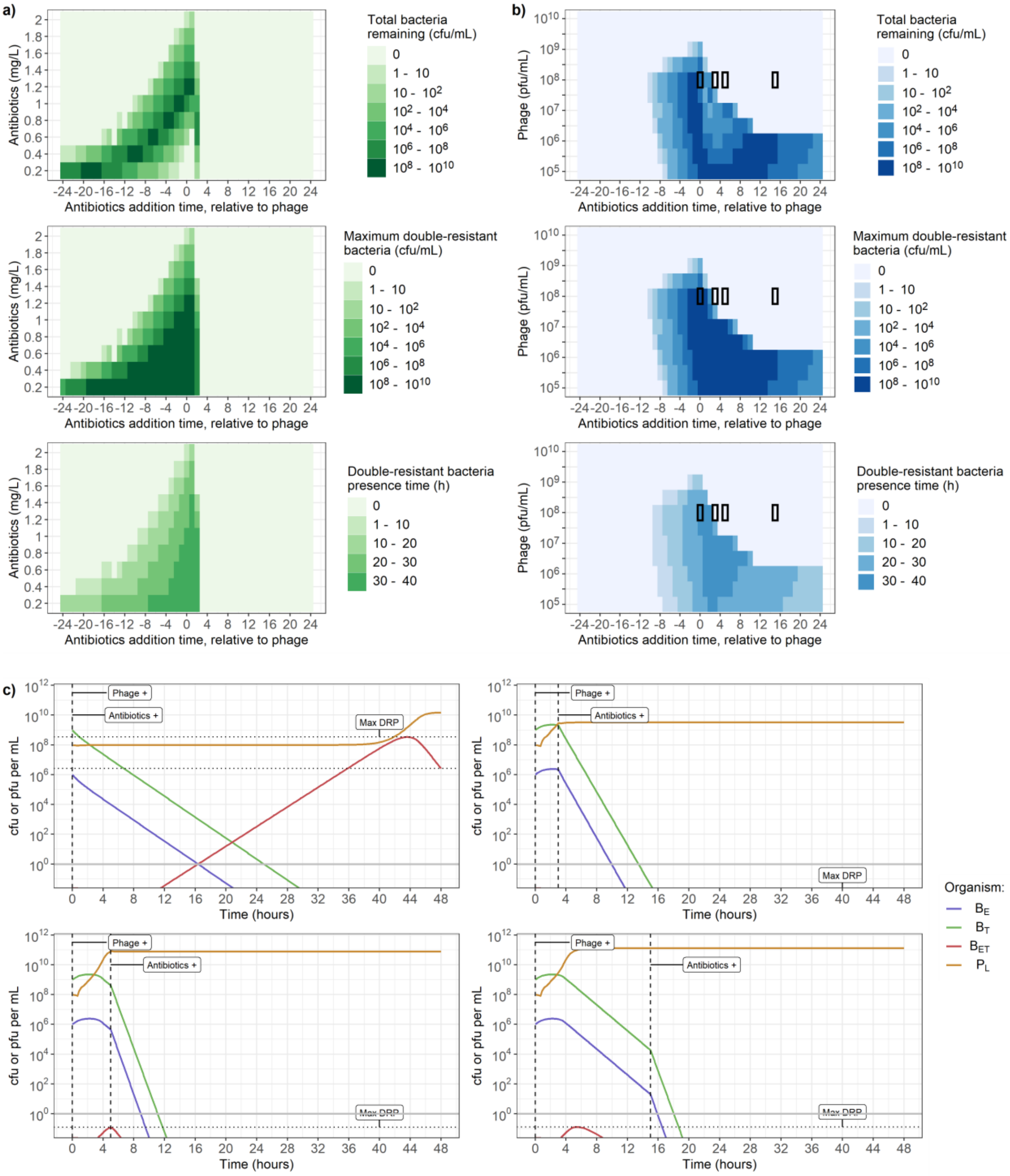


**Figure S5: Effect of varying antibiotic and phage timing and concentration when the erythromycin-resistant bacterial strain ($B_E$) is in minority ($10^6$ cfu/mL). a-b) Varying timing (x-axis) and dose of antibiotic and phage (y-axis) affects total bacterial count after 48h (top), maximum concentration of double-resistant bacteria ($B_{ET}$) (middle), and time when the concentration of $B_{ET}$ is greater than 1 colony-forming unit (cfu) per mL (bottom). a)** Adding $10^8$ plaque-forming units (pfu) per mL of phage, and between 0.2 and 2.2 mg/L of both erythromycin and tetracycline. **b)** Adding 1 mg/L of both erythromycin and tetracycline, and between $10^5$ and $10^{10}$ pfu/mL of phage. The x-axis indicates the time when antibiotics were added, relative to when phage were added. For example, the value "4"

indicates that phage were present at the start of the simulation, and antibiotics were introduced 4h later. The segments with black borders correspond to the dynamics shown in c). **c) Phage and bacteria dynamics over 48h for 4 conditions taken from panel b.** In all 4 conditions, indicated by the black rectangles, phage are initially present at a concentration of $10^8$ pfu/mL, while erythromycin and tetracycline are both introduced at concentrations of 1 mg/L after either 0h, 3h, 5h or 15h, stated on the plots, with the timing indicated by the vertical dashed lines. Horizontal dotted lines indicate bacteria remaining after 48h (corresponding to the top row of a-b) and maximum double-resistant bacteria ($B_{ET}$) concentration (middle row of a-b). Solid line indicates the detection threshold of 1 cfu or pfu/mL.

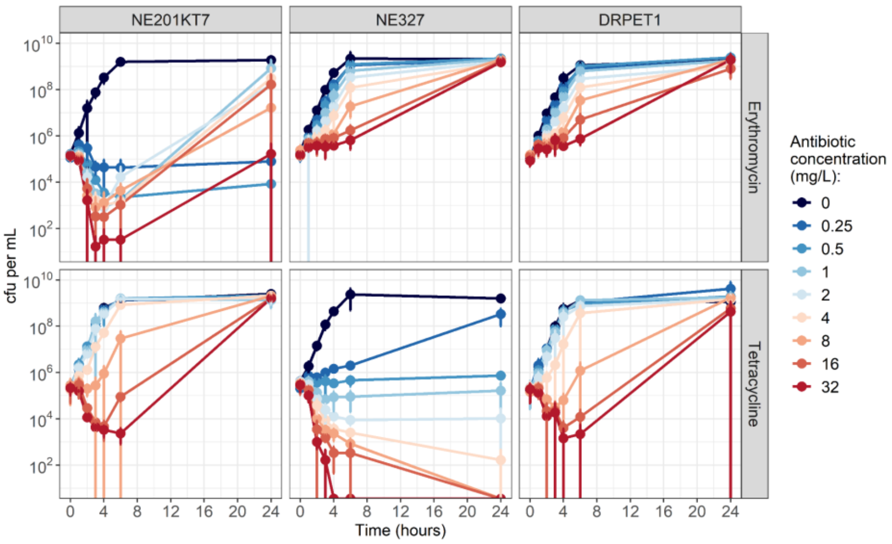


**Figure S6: Growth curves of NE201KT7 (tetracycline-resistant, left), NE327 (erythromycin-resistant, middle) and DRPET1 (double-resistant, right), exposed to varying concentrations of erythromycin (top) or tetracycline (bottom).** The minimum inhibitory concentration values for bacteria at 24h were identical to the ones for stock bacteria, suggesting that antibiotic decay rather than acquired resistance is responsible for the increase in bacteria numbers after 24h. Error error bars indicate mean +/- standard deviation, from 3 replicates. cfu: colony-forming units. Note that cfu per mL are shown on a log-scale.

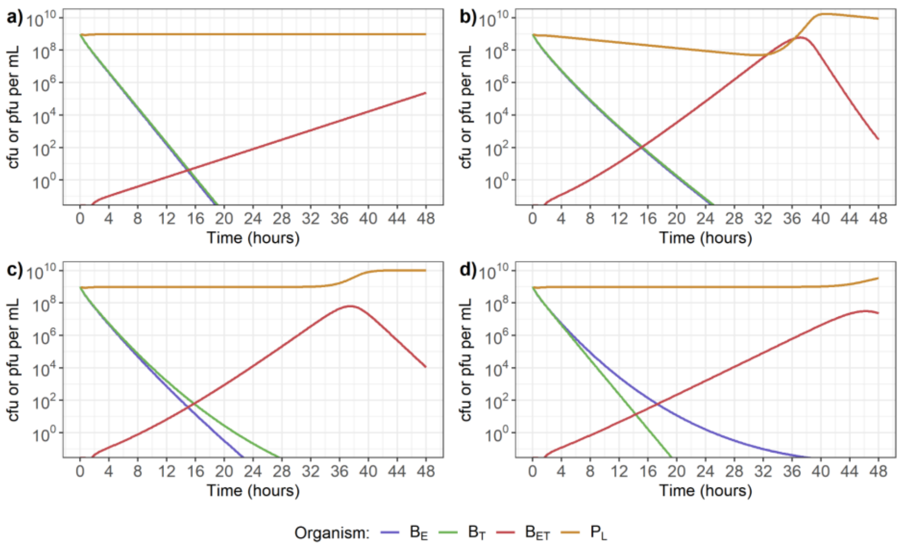


**Figure S7: Impact of phage and antibiotic decay on phage and bacteria dynamics over 48h.** The conditions shown are: no decay (a), phage decay (b), erythromycin decay (c), and tetracycline decay (d). In all 4 conditions, phage and antibiotics (erythromycin and tetracycline) are initially present at concentrations of $10^9$ pfu/mL and 1 mg/L respectively. Rates of decay are set to either 0 or 0.1 per hour.
